# Broad Spectrum Antimicrobial Potential of *Crassula ovata*

**DOI:** 10.1101/2022.01.20.477177

**Authors:** Troy Puga, Josh Schafer, Alexander Buffalo, Prince N. Agbedanu

**Affiliations:** Kansas City University School of Medicine, Kansas City, USA; University of Kansas School of Medicine, Kansas City, USA; Hutchinson Community College, Newton, Kansas, USA; Friends University, Department of Health Sciences, Wichita, USA

## Abstract

This study investigates the antimicrobial properties of *Crassula ovata* (*C. ovata*) against both gram-positive and gram-negative bacteria. About 1-3g samples of *C. Ovata* leaf samples were extracted with 95 % ethanol and the extract infused into paper discs by soaking and drying. The dried discs were screened against various strains of bacteria and the antimicrobial effects of the infused agents determined by measuring zones of inhibition due to the agents infused into the discs. By the zones of inhibition, *C. ovata* showed antimicrobial activity against the following gram-negative bacteria: *E. coli* (14 mm mean zone of clearing), *P. vulgaris* (13 mm mean zone of clearing), *E. cloacae* (16 mm mean zone of clearing), and *K. pneumoniae* (13 mm mean zone of clearing). *C. ovata* showed antimicrobial activity against the following gram-positive bacteria: *S. aureus* (22 mm mean zone of clearing) and *S. agalactiae* (8 mm mean zone of clearing). *C. ovata* did not show antimicrobial activity against *S. pyogenes*. By its antimicrobial activity against gram-positive and gram-negative bacteria, *C. ovata* displayed broad spectrum antimicrobial activity against *E. coli, S. aureus, S. agalactiae, P. vulgaris, E. cloacae, and K. pneumoniae. C. ovata* may have the potential to serve as a broad-spectrum antimicrobial in the future. Further testing should be done to investigate the toxicities and side effects of *C. ovata*. Further testing must also be done with *C. ovata* against drug-resistant strains of bacteria.

## INTRODUCTION

Many plants across the world have been used for medicinal purposes due to their antimicrobial properties. Plants in the modern era have become the source for the latest in antimicrobial development. *Crassula ovata*, more commonly known as Jade plant, is a native of South Africa (https://libguides.nybg.org/crassula). Many people in North America also keep *C. ovata* as a house plant (https://libguides.nybg.org/crassula). Historically, *Crassula ovata* has been used medicinally in both South Africa and China [2]. This experiment tested the following gram-negative bacteria: *E. coli, P. vulgaris, E. cloacae, and K. pneumoniae*. This experiment tested the *C. ovata* against the following gram-positive bacteria: *S. aureus, S. agalactiae, and S. pyogenes*.

Previously, *Crassula ovata* has been shown to have antimicrobial activity against *Eschericia coli* [2]. *E. coli* is a gram-negative rod that exists as part of the normal enteric flora [3]. It is a common source of both genitourinary and gastrointestinal tract infections, and is a common cause of watery and bloody diarrheal illnesses [3]. *E. coli* O157:H7 is a particularly lethal strain, known to cause Hemolytic Uremic Syndrome in children and the elderly [3]. *E. coli* has been proven to a pathogenic and potentially lethal microbe, which leads us to the issue that certain *E. coli* strains may have genes that confer antibiotic resistance [3]. *Proteus vulgaris* is another example of a gram-negative bacteria that has the potential to increase morbidity and mortality in immunocompromised individuals. *Proteus spp*. have shown the ability to confer some drug resistance [4]. *Klebsiella pneumoniae* (*K. pneumoniae*) is a gram-negative encapsulated bacterium that causes pneumonia, along with lung abscesses [5]. *K. pneumoniae* is a common source of nosocomial infection and has also been shown to gain antibiotic resistance [6].

*Enterobacter cloacae* (*E. cloacae*) is a gram-negative rod and multi-resistant bacteria that has been associated with large hospital outbreaks of gastrointestinal infection [7]. *Staphylococcus aureus* (*S. aureus*) is a gram-positive bacterium that is found in the normal flora of skin and mucous membranes and is a cause of skin abscesses, infective endocarditis, food poisoning, toxic-shock syndrome, and osteomyelitis [8].

Finding medications that have potent antibiotic activity against *Methicillin-resistant Staphylococcus aureus* (MRSA) is becoming increasingly difficult to come by, due to its extreme resistance to many antibiotics [8]. *Streptococcus agalactiae* (Group B Streptococcus) is a gram-positive coccus that is particularly dangerous to high risk groups such as neonates, pregnant women, and the elderly because of its ability to cause sepsis and shock [9]. Raabe and Shane, 2019). Some strains of *Streptococcus agalactiae* have shown resistance to penicillin and non-beta lactam antibiotics [9]. *Streptococcus pyogenes* (Group A Streptococcus) can cause pharyngitis, necrotizing fasciitis, and skin infections, along with long-term sequelae, such as post-infectious rheumatic fever and post-streptococcal glomerulonephritis [10].

## RESULTS

*Crassula ovata* showed antimicrobial activity against the following gram-negative bacteria: *E. coli* (14 mm mean zone of clearing), *P. vulgaris* (13 mm mean zone of clearing), *E. cloacae* (16 mm mean zone of clearing), and *K. pneumoniae* (13 mm mean zone of clearing). *Crassula ovata* showed antimicrobial activity against the following gram-positive bacteria: *S. aureus* (22 mm mean zone of clearing) and *S. agalactiae* (8 mm mean zone of clearing). *Crassula ovata* did not show antimicrobial activity against *S. pyogenes*.

## DISCUSSION

According to our experiments, *Crassula ovata* has shown broad-spectrum antibacterial activity due to its effectiveness against both gram-positive and gram-negative organisms (Table 1 and Table 2), which suggests that the extract of *Crassula ovata* could potentially be developed into a broad-spectrum antimicrobial agent. Further testing must be done against drug resistant strains of bacteria to test its full efficacy. Further testing also must be done in vivo to test for potential toxicities and side effects of *Crassula ovata*. Although the zone of inhibition measured for C. Ovata clearing of *S. aureus* (Figure 2) was greater in magnitude compared to that measured due to *C. ovata* clearing of *E. cloacae, E. coli*, and *K. pneumoniae* (Figure 2), it can only be interpreted as increased susceptibility but not increased potency [11]. The actual potency of *C. Ovata* against all strains has to be determined by in vivo treatments [11]. We welcome fellow scientists to conduct in vivo studies targeting these strains of bacteria in vivo and monitoring effects on the bacteria. Having demonstrated antibacterial activity against both gram-positive and gram-negative bacterial strains investigated, we conclude that *Crassula ovata* displayed broad spectrum antibacterial activity against *E. coli, S. aureus, S. agalactiae, P. vulgaris, E. cloacae, and K. pneumoniae*. We believe *C. ovata* may have the potential to serve as a broad-spectrum antimicrobial in the future.

**Table 1:** Gram-positive bacteria mean zone of clearing (in mm) against a blank disk and *Crassula ovata*.

|  | *S. aureus*<br>(clearing in mm) | *S. agalactiae*<br>clearing in mm) | *S. pyogenes*<br>(clearing in mm) |
| --- | --- | --- | --- |
| Blank Disk | 0 mm | 0 mm | 0 mm |
| *Crassula ovata* | 22 mm | 8 mm | 0 mm |

**Table 2:** Gram-negative bacteria mean zone of clearing (in mm) against a blank disk and *Crassula ovata*.

|  | *E. coli*<br>(clearing in mm) | *P. vulgaris*<br>(clearing in mm) | *E. cloacae*<br>(clearing in mm) | *K. pneumoniae*<br>(clearing in mm) |
| --- | --- | --- | --- | --- |
| Blank Disk | 0 mm | 0 mm | 0 mm | 0 mm |
| *Crassula ovata* | 14 mm | 13 mm | 16 mm | 13 mm |

**Figure 1:**
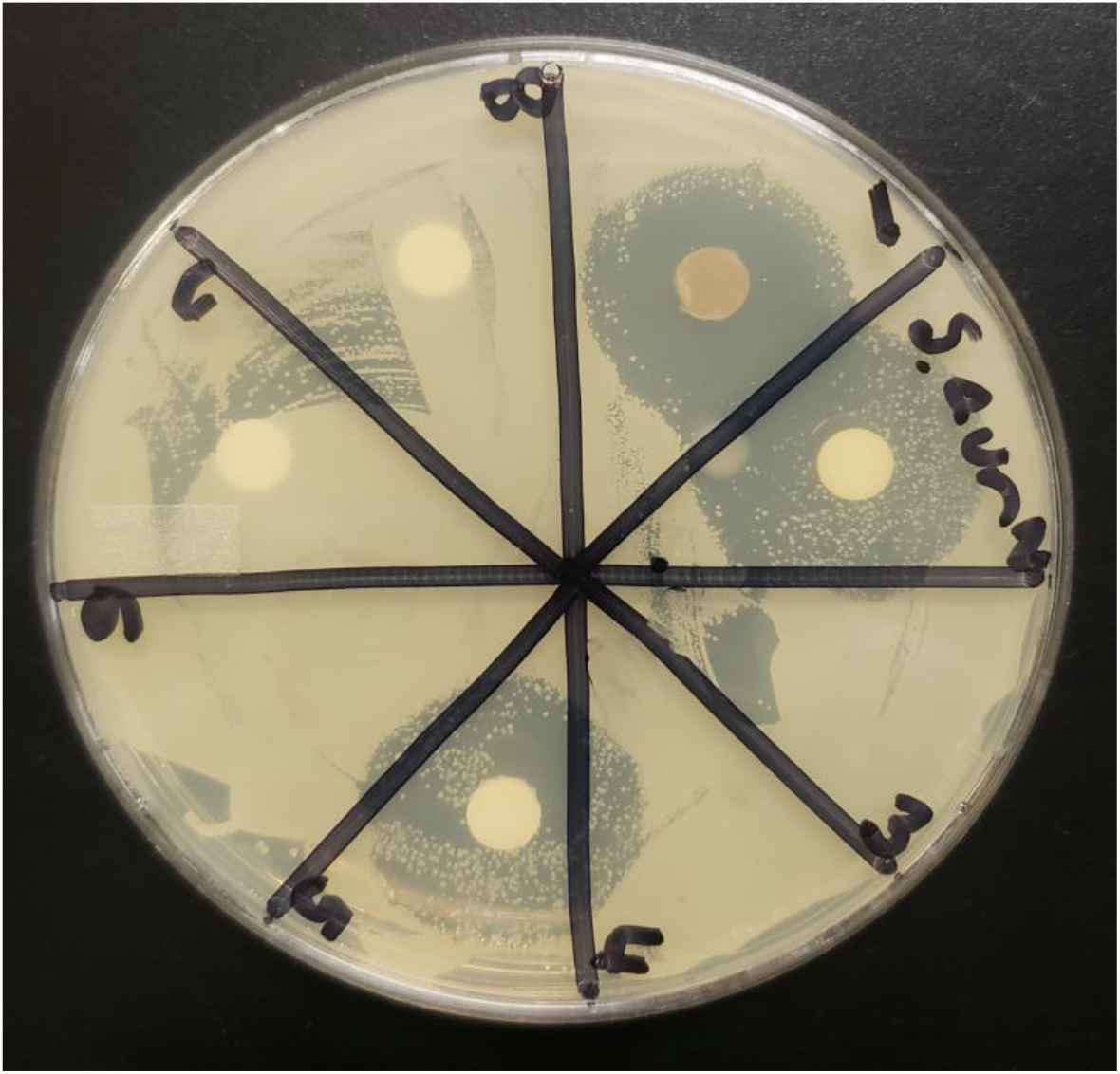
*Crassula ovata* (Position 1) and blank disk (Position 8); clearing against *S. aureus*. Zone of inhibition showing as a clear ring around disk, measured (diameter of zone in mm) and recorded in table 1.

**Figure 2:**
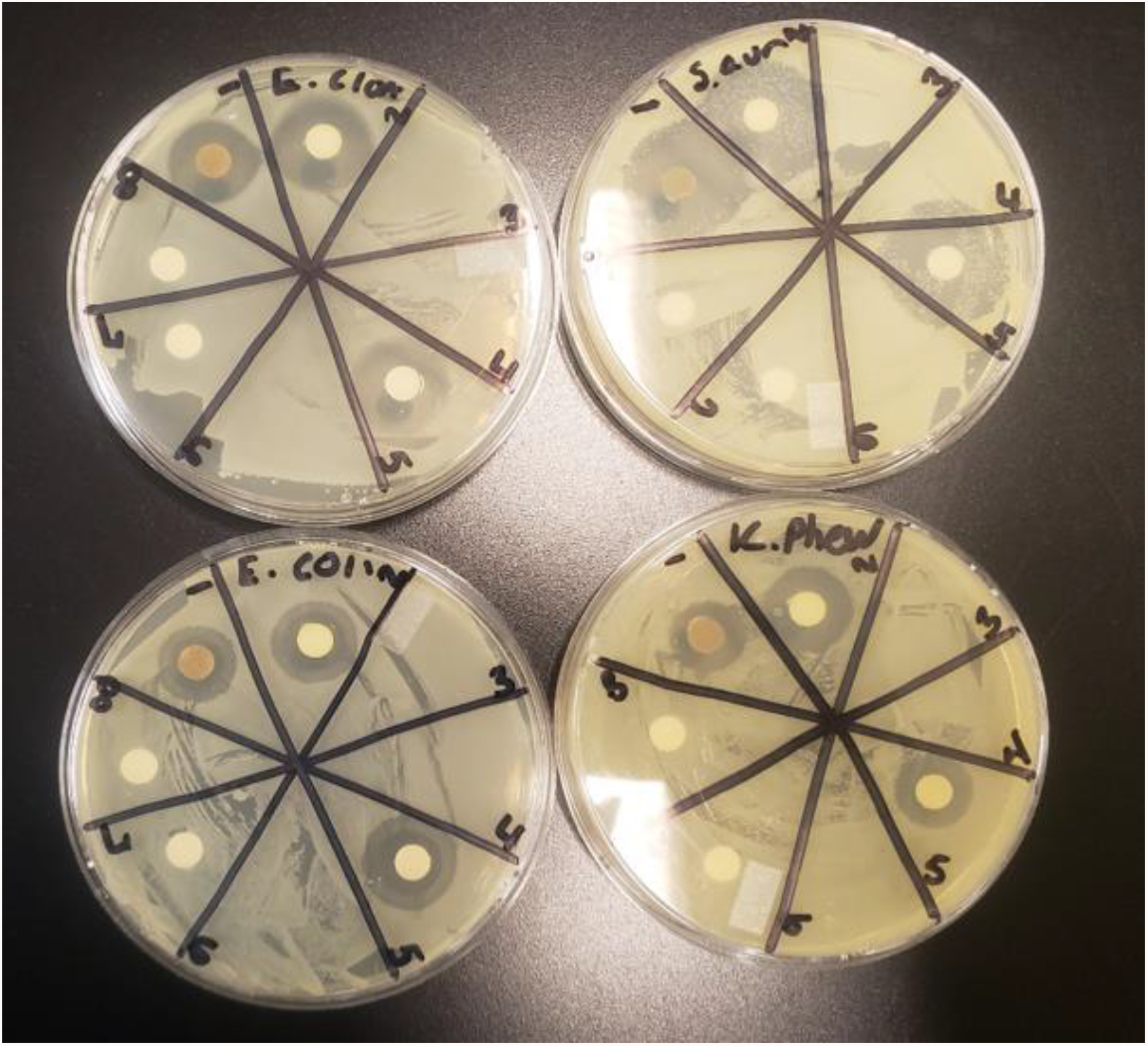
*Crassula ovata* (Position 1, all plates) and blank disk (Position 8, all plates) against *S. aureus, E. cloacae, E. coli, and K. pneumoniae*. Zones of inhibition showing as clear ring around disk, measured (diameter of zone in mm) and recorded in table 1 and 2.

## MATERIALS AND METHODS

### Sample extraction and disc preparation

1-3g samples of *C. Ovata* were homogenized using mortar and pestle to achieve a medium-fine paste. 3-4mls of 95% ethanol was added to the paste and swirled for 5 minutes to extract. The homogenate/ethanol mixture was filtered through a thin filter paper and the filtrate was collected into a 50 ml beaker. Blank disks were soaked into the filtrate and allowed to sit for 10 minutes.

The disks were removed using tweezers and allowed to dry on blotting paper. Other blank discs were soaked in 95% ethanol and allowed to sit for 10 minutes. The discs were removed and allowed to dry on a blotting paper for 30 minutes. During this wait time, bacterial plates were prepared from previously scaled bacterial stocks of the bacterial strains.

### Scaling up of bacterial from glycerol stocks

Bacterial stocks made from single colonies (500 microliters) were scaled-up in 1ml of Luria Bethani broth by shaking at 145 rpm and 37 degrees Celsius for 20 hours. A 100ul volume of the scaled-up stock was resuspended in 8ml of 1% saline solution.

### Preparing plates

Mueller-Hinton agar plates were prepared by dissolving 38g of nutrient agar in 1 liter of water. The mixture was autoclaved for 15 minutes at a temperature of 121 C. The media was then cooled and plates were poured, using approximately 20 ml volumes of the molten/autoclaved agar and allowed to set.

### Plating

100 microliters of the saline bacterial suspension were spread on Mueller Hinton plates. The discs with extracts (previously bathed in extracts) were placed on the plates; 4-7 discs/ per plate along with a blank disc (with no active ingredient impregnated, as negative control) or blank disc infused with the solvent used for plant extraction (control for solvent effect). The plates were incubated at 37 degrees Celsius for 24-48 hours and monitored for zones of inhibition.

## Acknowledgments

We thank the VPAA’s office and the Chair of STEM, Dr. Nora Strasser, of Friends University, for their support of both undergraduate and graduate research.

